# Global Phylogenomic Analysis and Genome Homology Reveals New rare Phylogenetic groups, “Synonymous” Species, and the Need to Simplify Nomenclature and Pathogens Identification in the *Bacillus cereus* Group

**DOI:** 10.1101/2025.03.11.642625

**Authors:** MH Guinebretiere, O Couvert, Y Le Marc, F Postollec

## Abstract

Bacteria in the *Bacillus cereus* Group *sensu lato* (Bc Group sl) may have serious detrimental effects on the health and safety of numerous foods. Over the past decade, phylogenetic studies have shifted toward the delineation of an increasing number of closely related species. For an improved understanding, this study attempts to fill existing gaps in the data to build a complete, robust and useful phylogenetic structure that contains all 29 currently described species.

The obtained phylogenetic structure features all of the known major phylogenetic groups (designated I to VII) and their subgroups, but also highlights the existence of seven new rare groups or sub-groups. The positioning of the 29 species revealed that some are synonymous, which casts doubt on the accuracy of their genomic delimitation. Four species matched with the previously described sub-groups III-5, III-9 and III-12, which are specifically known for their virulence potential. *B. thuringiensis* strains were dispersed throughout the majors groups II to VI, demonstrating that this species is not a genomic species restricted to group IV but should be instead considered a variety largely spread into the Bc Group sl. A similar pattern was observed for *B. mycoides* in 10 phylogenetic groups/species.

These results highlight the need for a simplified nomenclature of the Bc Group sl, which should be developed through a collaboration between relevant stakeholders. In the meantime, an updated version of the *panC* database is proposed, named CereusID, that integrates data from the present work and enables the identification of all phylogenetic entities within Bc Group sl: https://toolcereusid.shinyapps.io/cereusid/.

## 1. Introduction

Bacterial strains in the *Bacillus cereus* Group *sensu lato* (Bc Group sl) present major challenges to the food industry. They contaminate a broad range of diverse foods, cause diarrheal and/or emetic syndromes, and play an important role in food spoilage. Furthermore, an incomplete understanding of their taxonomy, which in recent years has become increasingly complex, hampers investigation of their accurate identification in foods or the environment.

In 2008, Guinebretiere et al. [1] described a phylogenetic structure for this bacterial Group that was composed of seven major phylogenetic groups (I to VII); these findings were based on a robust analysis of restriction-fAFLP features and *rrs-its* sequences of 435 independent bacterial strains selected around the world. Each of the seven major phylogenetic groups was associated with an identifiable ecological niche as well as a specific range of growth temperature, a trait that is often important in bacterial taxonomy. The ‘thermotypes’ thus defined were then further linked with different risks of food poisoning [2]. In addition to these well-known phylogenetic groups, 14 sub-groups were described in 2008 [1] and further characterized in [2]; each sub-group was also associated with a specific risk of food poisoning, but these have received scant attention by the scientific community. In these early studies, the most important phenotypic features were more often group-dependant than species-dependant [2].

Until 2010, the phylogenetic structure for the Bc Group sl contained six species (*B. cereus, B. thuringiensis, B. anthracis, B. mycoides, B. weihenstephanensis*, and *B. pseudomycoides*). Among these, *B. thuringiensis* is of great importance because it exhibits highly effective insecticidal properties and is employed as a bio-pesticide around the world. For many years, it was considered a separate species from *B. cereus*, but is now recognized to be affiliated to the Bc Group sl. However, its genetic and phenotypic diversity, as well as its relationship with other species in this Group, remain unclear for many authors still nowadays.

Since 2010, numerous new species have been described, adding to the richness of the Bc Group sl but also increasing its taxonomic complexity. Indeed, 24 closely related species have been described in the last decade [3-14], reaching 29 species today. During their description, the phylogenetic positions of these species within the entire Group have been partially or not accurately resolved. This also resulted in confusion or gaps in discriminative phenotypic characters and virulence potential. Finally, within this ambiguous taxonomy, *B. thuringiensis* oscillates between different phylogenetic positions depending on the study in question: some authors place it within phylogenetic group IV [15] while others consider it to be widely spread throughout the Bc Group sl [1].

The aim of this work was to obtain the global, complete, and robust phylogenetic structure for the Bc Group sl, based on data from 1000 genomes, and that allows to clarify (i) position of all constitutive phylogenetic groups and sub-groups, including new entities, (ii) position of the 29 current species into that phylogenetic structure, with their phenotypic and virulence specificities. It also present the resulting useful application for strain identification, and their virulence potential.

## 2. Material and Methods

### 2.1 Phylogenomic Analysis

The complete phylogenetic structure of the Bc Group sl was established based on the evolutionary history of 1000 genomes; the large volume of data used in this analysis increased the robustness of branches and thus our confidence in the results.

Phylogenomic tree reconstruction was performed using the PATRIC web-based resource (PAThosystems Resource Integration Center) [16, 17]. This process reconstructs high-quality phylogenetic trees using an automated pipeline that includes maximum likelihood (ML) tree inference, alignment trimming, and evolutionary model testing [18-24]. To obtain the complete phylogenetic structure, 30 trees were reconstructed through a process composed of several steps, described below.

#### 2.1.1. Genomes databases

The first dataset constituted 45 reference genomes, which were selected as representatives of all known diversity in the Bc Group sl (i.e., phylogenetic groups I to VII, their sub-groups, and species), from PATRIC database [17].

The second dataset comprised 955 genomes randomly selected from the PATRIC database [17], from the set of all known strains with phylogenetic or species affiliation to Bc Group sl (as determined by PATRIC pre-analysis). For groups or species that were underrepresented, we included all strains available.

Three private genomes of *B. thuringiensis* were included, which were representatives of the main active strains used as bio-pesticides in Europe.

All genomes included in this study are listed in Supplementary data **S1**.

#### 2.1.2. Phylogenetic affiliation of 955 genomes based on comparison to the 45 reference genomes

All of the 955 selected genomes were compared to the 45 reference genomes in randomly selected sets of 55; the analysis was reiterated until all 955 genomes had been used. Based on this, a total of 18 ML trees were reconstructed using *B. manliponensis* JCM 15802 as the outgroup. From those 18 trees, 55 sub-trees were generated to accurately examine specific branches. All 955 genomes were affiliated to known or unknown phylogenetic groups (supplementary data S1).

#### 2.1.3. Phylogenetic analysis of genomes, toward building of 14 detailed trees and the final Global tree

We eliminated 42 genomes that were either of poor quality (high number of contigs, low depth, errors, defect checking) or too similar to other genomes (clonal genomes, duplicate genomes). The remaining 958 genomes (including reference strains) were assorted by their phylogenetic affiliation, and each resulting set of genomes was compared on a single detailed tree at the sub-group (or finer) level. This step resulted in 12 detailed trees and two sub-trees.

Finally, 100 genomes were selected from the 14 detailed trees as representatives of all the phylogenetic diversity present. These were used to reconstruct a final, global tree exhibiting the entire phylogenetic structure of Bc Group sl, and rooted on *B. manliponensis* JCM 15802.

### 2.2 ANI calculation and percentage of conserved DNA

ANI value and the percentage of conserved DNA [25] were calculated for all pairs of type-strains and group-representatives of the Bc Group sl. Both parameters were calculated using the ANI script (kindly provided by K. T. Konstantinidis, Michigan State University), at 85% similarity level and at minimum 70% coverage. The final data are means of reciprocal values for each pair of strains.

## 3. Results and Discussion

### 3.1. Toward a more comprehensive phylogeny of the entire Bc Group sl

The present phylogenomic analysis, including 1000 genomes from members of the Bc Group sl, represents the most exhaustive effort to date to capture the diversity of this species complex and to robustly delineate all phylogenetic entities contained therein. As part of this work, we constructed a global tree (Fig. 1) along with 14 detailed sub-trees (available from the corresponding author) on which all known phylogenetic groups (I to VII), their sub-groups [1, 2], new entities, and all 29 described species are depicted in relation to each other.

**Fig. 1.**
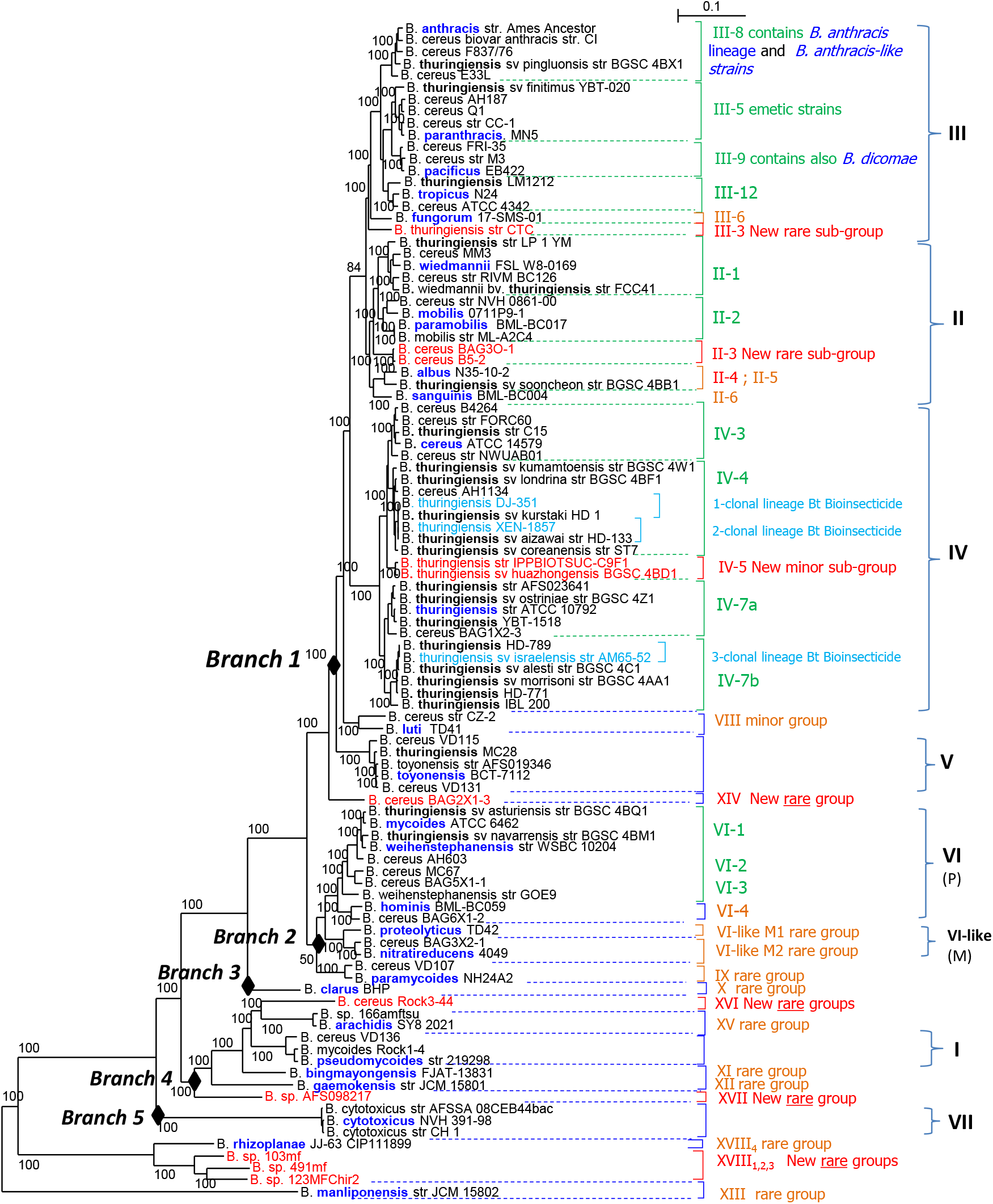
Global Phylogenetic Tree of the *B. cereus* Group. MLTree based on the core genome and rooted on *B. manliponensis. The 100 genomes were selected as representatives, based on the present phylogenomic study including 1000 genomes. Species names are originating names, except for type-strains*. **I** to **VII** : major groups according to Guinebretiere et al., 2008 ; In GREEN : sub-groups defined by Guinebretiere et al. 2008, 2010 ; In ORANGE: groups and sub-groups defined in the present study ; In RED : New groups or sub-groups find in the present study. (P): Psychrotolerent ; (M): Mesophilic

As shown in Figure 1 and Table 1, the high majority of the obtained phylogenetic structure is composed of the seven major phylogenetic groups (I to VII) and their sub-groups, which were described in 2008 and 2010 [1, 2]. However, there are also new **rare** or **minor** groups, mainly positioned closely related to group VI or in the lower part of the global tree (VI-likeM1, VI-likeM2; VIII to XVIII). In the middle and upper part of the global tree (Fig. 1, Table 1), there are new minor or rare sub-groups within the known groups (III-3, III-6, II-3, II-4, II-5, II-6, IV-5, VI-4). Among all these, six rare groups (XIV, XVI, XVII, XVIII-1,2,3) and four rare to minor sub-groups (III-3, II-3, II-4, IV-5) do not match with any known group, sub-group, or species, and thus might be considered as-yet-undescribed phylogenetic entities (Table 1).

**Table 1.**
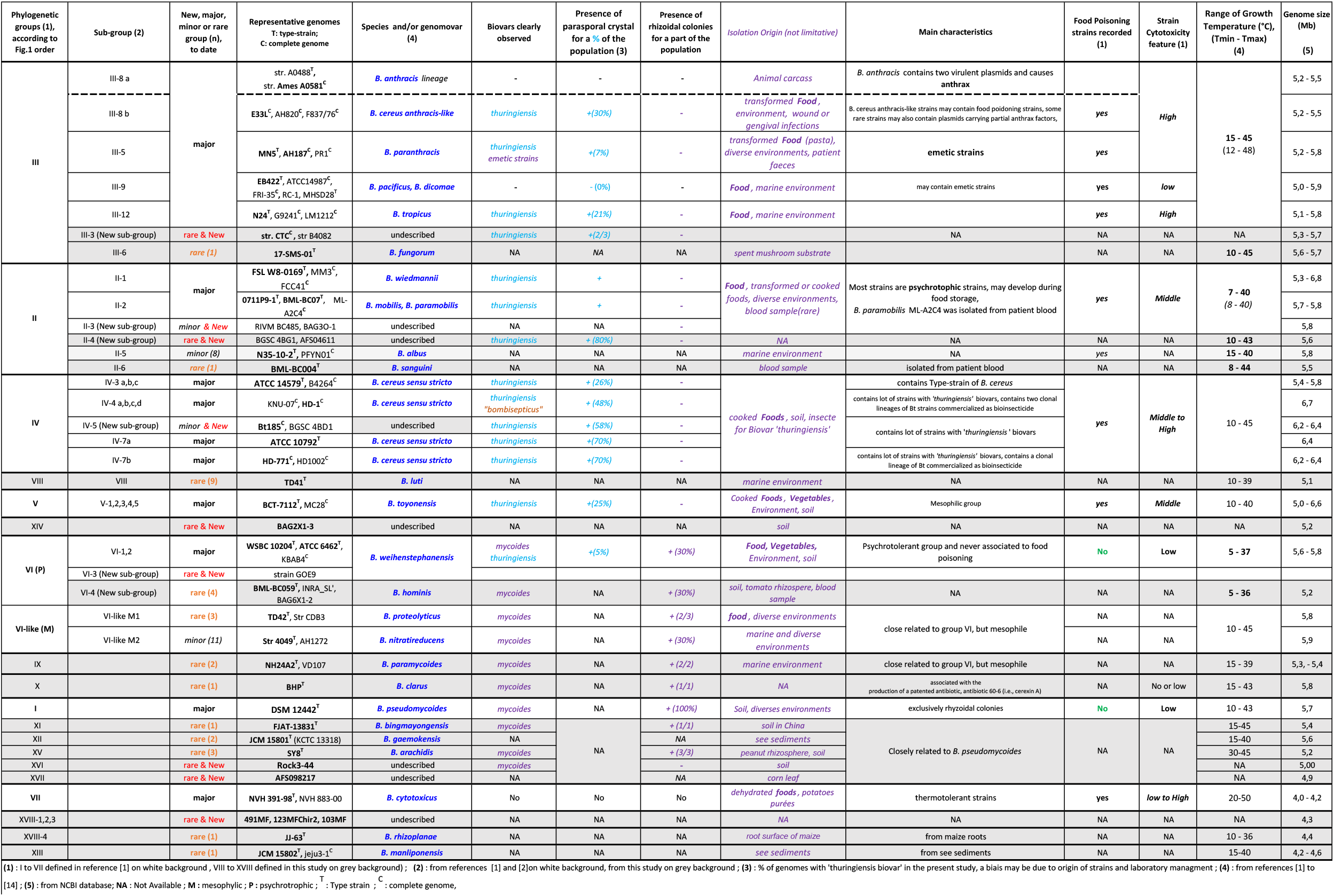
Mains characteristics of Phylogenetic entities and Species found into the *B. cereus* Group.

This work confirms the previous assignments of *B. cytotoxicus, B. pseudomycoides, B. weihenstephanensis/B. mycoides* to groups VII, I, and VI, respectively [1, 2]. Likewise, *B. paranthracis, B. pacificus, B. dicomae*, and *B. tropicus* were again found to be genomically affiliated with the well-described phylogenetic group III, in sub-groups III-5, III-9, III-9 and III-12, respectively [2]. *B. fungorum* formed a new sub-group in group III (named III-6). *B. wiedmanii, B. mobilis, B. paramobilis, B. sanguini*, and *B. albus* were assigned to group II, *B. toyonensis* to group V, and *B. hominis* to group VI (sub-group VI-5). All the remaining species (*B. luti, B. nitratireducens, B. proteolyticus, B. paramycoides, B. clarus, B. arachidis, “B. bingmayongensis”, “B. gaemokensis”, “B. manliponensis”*, and *B. rhizoplanae*) matched with new and **very rare groups** : VIII, VI-LikeM1, VI-LikeM2, IX, X, XV, XI, XII, XIII, and XVIII-4, respectively (Fig. 1, Table-1). For most of these rare groups, there is not any knowledge on their virulence potential.

### 3.2 A specific focus on B. thuringiensis distribution throughout the overall Bc Group sl

In the present study, *B. thuringiensis* genomes were largely represented by strains from the BGSC collection [https://bgsc.org/]. Contrary to what was previously proposed for this species [15], isolates of *B. thuringiensis* were not restricted to group IV (Fig.1, Table 1). Instead, these genomes could be found in almost every major phylogenetic groups I to VI, may be even in the lower part of the global tree (Fig.1, Table 1). Indeed, given this range, it is reasonable to question if we would have also found *B. thuringiensis* within other groups or species if they were not so poorly represented. This large phylogenetic diversity presumably explains the large diversity of insecticidal toxins described for this crystal-making bacterium, and definitively supports *B. thuringiensis* as a biovar/pathovar of few species/genomovars instead of a separate and well delimited genomovars. *B. thuringiensis* biovar was previously observed for *B. wiedmannii* [26]. This biovar/pathovar appears to occur in almost all the major phylogenetic groups and/or species within the Bc Group sl, and is defined by the production of insecticidal proteins that are easily observable as parasporal crystal(s) produced in the sporulation stage.

As shown in Fig. 1, the three main strains of *B. thuringiensis* that are used as Bt bioinsectide in Europe are restricted to three clonal lineages within the sub-groups IV-4 and IV-7b. Although the *B. thuringiensis ‘*biovar’ should in theory inherit all of the properties of the phylogenetic group and/or species to which it belongs (including virulence properties). However, these three highly host-specific strains have been widespread in the environment for a long time without any reported association to food poisoning, and presumably represent no or low risk. An interesting hypothesis to explain this would be the exclusive host specificity of *B. thuringiensis* concerning the insect, in other words, the safety of *B. thuringiensis* for humans. These findings are consistent with the reputation of the Bt bioinsecticide strains as well-delimited to three lineages and easy-to-characterize bacteria that can be analysed case by case for new formulations. This type of strains should be considered separately from the general character of the *B. thuringiensis* ‘pathovar/biovar’.

In the Bc Group sl, should we maintain *B. thuringiensis* as a distinct species or should it be classified as a biovar/pathovar of the Bc Group sl? In the event that the host specificity of *B. thuringiensis* makes it innocuous to humans, and considering the significant role it plays in biological control, it might be reasonable to keep this bacterium as a distinct species and maintain its nomenclature.

### 3.3 Species delimitation within Bc Group sl

#### 3.3.1. Synonymous species and/or groups

The present phylogenomic analysis highlights certain **species** that are potentially **synonymous**: *B. mobilis and paramobilis, B. paranthracis, pacificus and B. dicomae, B. proteolyticus and B. nitratireducens* (Fig. 1 and Table 2), although they may be distinguished at a sub-group level. This pattern can be noted particularly in groups II, III, IV and VI, with ANI values and conserved-DNA ≥ 96% and ≥ 70% respectively (Table 2). This situation is probably due to the fact that, in the last decade, an extraordinary number of species were independently described at the same time, so that different authors were not able to take all of them into account in phylogenetic analyses. In addition, use of 16S sequence analysis did not provide the discriminatory power to accurately distinguish species within these bacterial Groups, specifically for *B. cereus* branch N°1 (groups II to V, Fig. 1) and N°2 (groups VI and VI-like, Fig. 1), thus resulting in weak phylogenetic positions. Finally, it is likely that species described since 2013 have been delimited using different homology thresholds, depending on the *in silico* method of analysis selected by the authors and the parameters applied for these methods. For example, the ANI value (Average Nucleotide Identity), which has been widely used for Bc Group sl species, may vary according to parameters such as the percent identity threshold for calculation and the minimal length of matched DNA-fragments selected by authors [25]. In addition, the ANI method has been adapted by different web-based tools for different types of data (e.g., WGA contigs, genes, orthologous genes, preselected genes), which necessarily introduces variations. Indeed, when we re-analysed strains representatives of species and/or phylogenetic entities in a single set, using the same method, far fewer genomic species were defined (Table3, detailed calculations are available from the corresponding author). Values of ANI and conserved DNA calculated between representatives of groups II to V (Branch 1, Fig. 1; Table 3) were high, 91.4% to 96.0% and 61% to 80%, respectively. All these data indicate that groups II to V, and their two rare and satellite groups (VIII and XIV), should be considered sub-species of *B. cereus* branch 1. Branch 1 should itself be considered the *B. cereus* species sensu stricto, which would include 4 sub-species (entities II to V), *B. luti* (VIII), and the rare group XIV, along with 12 genomovars (*paranthracis, pacificus, dicomae, tropicus, fungorum, wiedmannii, mobilis, paramobilis, albus, sanguini, luti*, and *toyonensis*), as well as 1 biovar or “horizontal species” (*B. thuringiensis*). This would result in the following nomenclature for Branch 1, for example, for the current *B. tropicus* species: *B. cereus* subsp. III gv *tropicus*. This nomenclature should be discussed for each branch, in the future, by an international board composed of *B. cereus* experts.

**Table 2.**
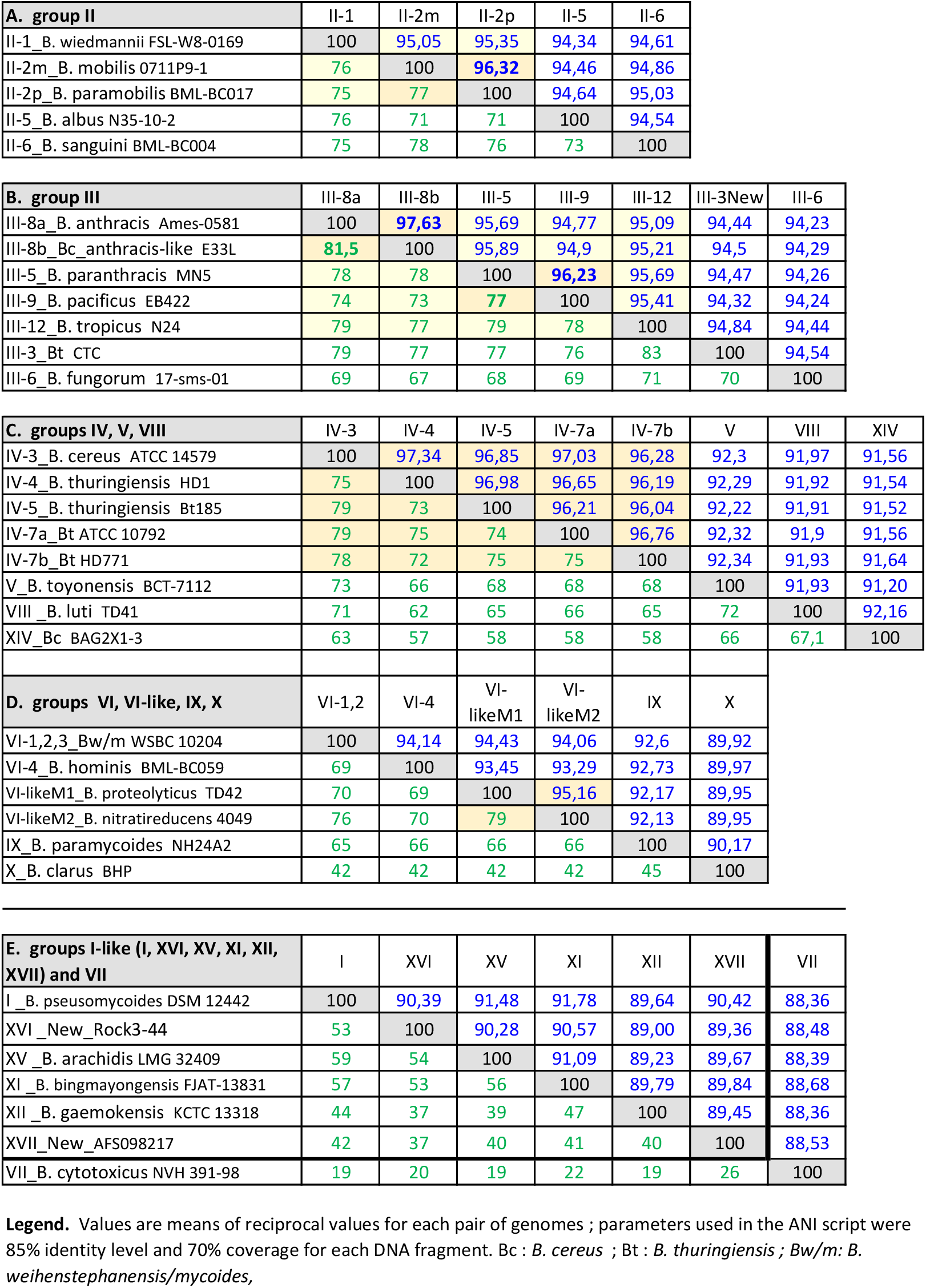
Intra-group ANI (in blue) and Conserved DNA (in green) between representatives genomes.

**Table 3.**
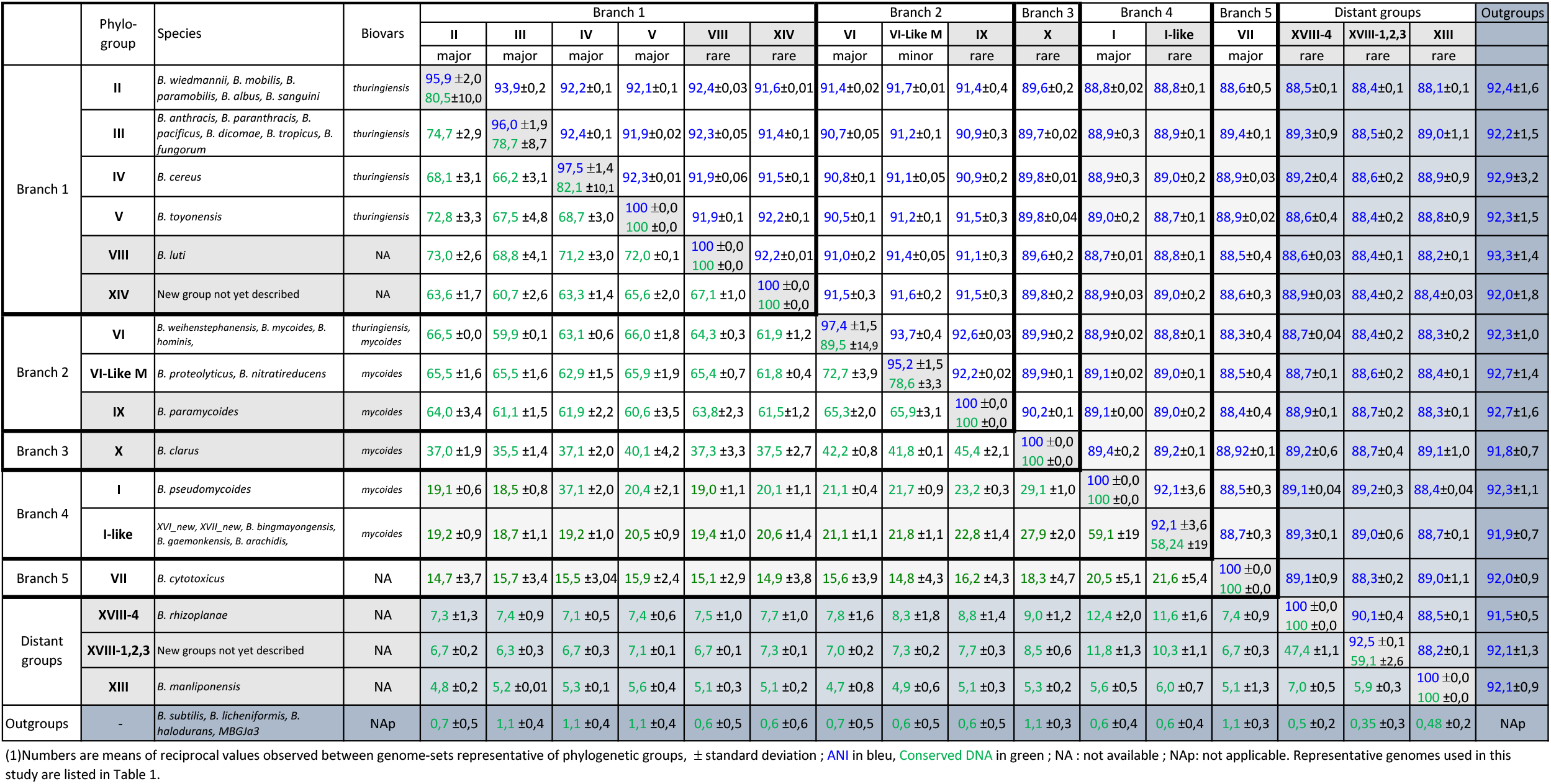
Inter-group ANI and Conserved DNA, between representative genomes ^(1)^

#### 3.3.2. The specific case of B. mycoides and B. weihenstephanensis within group VI

The name “*B. mycoides”* came from this bacterium’s singular ability to produce filamentous growth, which results in rhizoidal or “mycoidal” colonies on agar medium, first described by Flügge 1886, and aggregation in liquid culture [27, 29]. Recently, the name “*B. weihenstephanensis*”, which belongs to the same genomospecies as *B. mycoides* in group VI, was replaced by “*B. mycoides”* [29]. Therefore, numerous group VI strains that do **not** harbor “rhizoidal” characteristics were or will be inappropriately re-named “*B. mycoides*”. *B. weihenstephanensis* (non-rhizoidal form) is more economically important to the food industry than *B. mycoides* (rhizoidal form), as it occurs more frequently in food and is responsible for food spoilage at low temperatures; for this reason, it would be useful to maintain the distinction between the two forms. Even more important, if we look more closely at Figure 1 and Table 1, we find that *B. mycoides* (rhizoidal form) is in fact a subset of different species, including *B. weihenstephanensis* within group VI, and should thus be considered a phenotypic variant (biovar) that expresses an unusual rhizoidal phenotype. In this study, the rhizoidal phenotype occurred in group-species other than VI-*B. weihenstephanensis*, such as VI-likeM1-*B. proteolyticus*, VI-likeM2-*B. nitratireducens*, IX-*B. paramycoides*, X-*B. clarus*, XV-*B. arachidis*, XVI-new-rare-group, I-*B. pseudomycoides*, XI-B. *bingmayongensis*, XII-*B. gaemokensis* (Table 1). *B. pseudomycoides* is the only major group in which 100% of strains exhibit the rhizoidal phenotype. This definitively supports *B. mycoides* as a biovar of few species/genomovars.

#### 3.3.3. The specific case of Groups II & III

Some authors have suggested merging group II and group III together to create “*B. mosaicus*” [15]. We propose to keep these two phylogenetic groups separate for the following reasons. Group III is clearly genomically separate from group II in the present phylogenomic study (Fig. 1), as well as in previous studies [1, 2, 30, 31], even if the *panC* sequence of a part of sub-group III-12 is highly similar to that of group II and bring confusion. ANI and conserved DNA were high between groups II and III (93 to 94 and 69% to 74% respectively), but high values are also observed between groups II to V (see above). Additionally, the thermal resistance and growth temperature ranges of these two groups are quite different [1, 2], with a shift toward low temperatures for group II compare to group III (Table 1). These characters are important not only for taxonomic purposes but also for industrial processes and food storage. Finally, the risk of food poisoning associated with group III is higher than with group II [2]. We thus encourage keeping group II and group III separate.

#### 3.3.4. Other observations on species delimitation within Bc Group sl

- Sub-group III-8 contains bacteria with important effects on health: *B. anthracis* and *B. anthracis*-like strains are responsible for, respectively, anthrax and serious illness (including superinfection of the lungs, periodontitis, wound infection, and food poisoning) [Table 1]. Within sub-group III-8b, the *B. cereus* strains closely related to *B. anthracis* would merit the name “paranthracis”. Unfortunately, this name has already been attributed to another sub-group in group III, subgroup III-5, which harbours emetic strains. Sub-groups III-5 and III-9 are closely related, and constitute the two main sub-groups containing emetic strains [Table-1].
- Representativeness of species: certain species are represented by only a few strains (less than three) and are thus considered as rare [Table 1]. It is only logical to ask questions regarding the significance and reliability of properties associated with them. Furthermore, there is always a risk of artefacts for species that are so poorly represented.
- Levels of ANI and Conserved DNA values observed in Table 3, between Branchs-1-to-5 and the distant groups XVIII-4 (*B. rhizoplanae*), XVIII-1,2,3 (new rare groups) and XIII (*B. manliponensis*), are very low, 88-89% for ANI and only 4 to 8% for Conserved DNA. Thus, affiliation of theses distant groups to the Bc Group sl should be also discussed by *B. cereus* experts and taxonomists.

### 3.4 Repercussions for food microbiology, and updating the panC database (now named CereusID, https://toolcereusid.shinyapps.io/cereusid/)

Prior to this analysis, 29 closely related species had been described within the Bc Group sl. Of these, numerous were revealed by this work to be fairly close relatives or even synonymous, majors groups II to V being associated below to the theoretical sub-species level. This greatly increases the taxonomic complexity of this bacterial group without any gain. Furthermore, after 2013, most published descriptions did not investigate the potential virulence of new species [Table 1]. This may create problems for both the food industry and health and safety organizations, who have faced difficulties in establishing a simple and coherent schema for microbiological analyses of the Bc Group sl.

In this study, 19 described species were found to be phylogenetically affiliated with one of the well-characterized phylogenetic groups (I to VII) and/or their sub-groups [Fig. 1, Table 1]. While most of the 29 species have not been assessed for their food poisoning risk, the major phylogenetic groups have [Table-1] [2]; and these phylogenetic assignments may therefore provide invaluable clues to the potential behavior of strains (ranges of growth temperature and potential virulence). Fortunately, the new groups identified by the present work, which have not yet been described, are rare (Table-1), and their scarcity and atypical sources suggest a low risk of occurrence in foods. For some, however, their proximity to known phylogenetic groups might give an indication of their level of risk.

The *panC* sequence remains particularly reliable for the identification of phylogenetic groups I to VII, and an online tool dedicated to this purpose has been available since 2010 [2]. As part of the present study, the *panC* database was updated to take into account all new data available from the present study, and the result was named CereusID. This exhaustive database includes the entire *panC* sequence from 287 genomes representing all phylogenetic entities found in the present work. Each sequence in the CereusID database was labelled with the group-affiliation of the genome from which it originates.

To improve the reliability of analyses, CereusID tool propose to users to match their isolates to existing phylogenetic groups (known and not yet described) instead of species. However, users also have the ability to identify the species to which their isolates belong, through both the global phylogenetic tree and a table summarizing the main known characteristics for each phylogenetic entity, including 28 species. *B. dicomae* [32] was identified to sub-group III-9 using CereusID tool. In 2025, the 29^eme^ species (*B. dicomae*) will be updated in the database and on the integrated tree.

In addition to groups I to VII and the new groups found here, the CereusID online database now allows the identification of isolates at the sub-group level, which may help in the detection of potential emetic strains (sub-groups III-5 and III-9) and *anthracis*-like strains (sub-group III-8b). It may also help to refine the identification of certain lineages such as those of *B. anthracis* (III-8a) and the three *B. thuringiensis* strains used as bioinsecticides in Europe.

The CereusID interface has undergone a complete redesign, with enriched user guidance through several “tabs” linked to the results table of the main page. Specifically, the web interface offers to users:

i. all recommendations for improving the reliability of phylogenetic assignments (“Preparation protocol” and “Sequence input”),
ii. information on identification methods and troubleshooting (“Identity” and “Troubleshooting”),
iii. known key tests that might improve or confirm the potential characteristics of isolates (“Additional tests” and “Characteristics”), especially if they originated from food,
iv. Positioning of species through a global phylogenetic tree on which users can position their isolates (“Phylogenetic tree”), and
v. detailed percent identity results compared to the entire database (“Identity tabs”).

We hope that this upgraded version will help members of the food industry, along with safety and health laboratories, to improve their microbiological identifications and to better evaluate the risk these bacteria pose for food products and health. The CereusID tool will remain flexible and will adapt as needed to changes in the taxonomy and nomenclature of the Bc Group sl.

## 4. Conclusion

As it currently stands, the large number of species in the Bc Group sl is unmanageable by many of the organizations and individuals who work with these bacteria, including members of the food industry, private quality-assurance laboratories, and public health laboratories. The problem is compounded by the fact that most of these species are highly similar, sometime synonymous, and/or have not been evaluated for their virulence potential. Furthermore, the most important phenotypic features of these bacteria are in fact more often group-dependant than species-dependant. The aim of this work was not to reassess the Bc Group sl, even if it provides key elements. It was to characterize its current state by defining the phylogenetic relationships between all the entities contained therein. It reveals several incoherencies and highlights the need for a re-evaluation of this species complex, as a simplified nomenclature, with the aim of producing a classification that is more useful for the microbiological analysis of foods and of clinical samples. Such work should be performed through a collaboration between all interested parties, including taxonomists, species description authors, food microbiologists, and health experts.

In the meantime, the updated phylogenetic structure presented here should help researchers to better understand the phylogenetic position of their isolates and manage the risks associated with them via the information available for the major and well-characterized phylogenetic groups and/or sub-groups. The data generated in the present work have been used to create a new *panC* database, CereusID, which should serve as a useful tool for exploring the large phylogenetic diversity of the Bc Group sl and the potential risk associated to new isolates.

## Supplementary data

**Supplementary data S1**. List of genomes analysed in this study and their resulting phylogenetic position.

## Data Availability Statement

All 18 positioning trees (N°1 to 18), all 14 detailed trees and sub-trees (N°19 to 30, 19-2, 19-3), and detailed Homology calculations (ANI, Conserved DNA) are available from the corresponding author by writing to. Particularly, the 14 detailed trees allow scientists to obtain the position at the sub-group level of several hundreds of genomes from international databases, if required for their own research.

## Funding and Acknowledgments

This study was funded by the French Ministry through the Casdar projet BtID (SA.40312-agreement N° 1528 BT ID), part of which came from self-funding from MICA division of INRAE. This work was also supported by UMT ACTIA TRANSISPORE (Joint Technological Unit - French Ministry responsible for Food).

For the three private genomes, we acknowledge the sequencing and bioinformatics expertise of the I2BC High-throughput sequencing facility, supported by France Génomique (funded by the French National Program “Investissement d’Avenir” ANR-10-INBS-09).

We also thank The Bacterial and Viral Bioinformatics Resource Center (BV-BRC). The BV-BRC including PATRIC Resources, has been funded in whole or in part with federal funds from the National Institute of Allergy and Infectious Diseases, National Institutes of Health, Department of Health and Human Services, under Contract No. 75N93019C00076 to the University of Chicago.

Projet design, Phylogenomic study and homology calculations: MHG; Conceptualization of CereusID online-tool: MHG, OC, YL and FP; panC Database: MHG; Interface design of CereusID, pages-content, data curation and resources: YL, MHG; Informatics’ program: OC, YL, MHG; writing-original draft: MHG; writing-review and editing: MHG, YL, OC, FP; Project Administration and funding acquisition: FP, MHG. All authors have read and agreed to the published version of the manuscript.

